# CircRNAs and their regulatory networks associated with the antioxidant enzymes and immune responses of Asian honey bee larvae to fungal infection

**DOI:** 10.1101/2023.07.19.549790

**Authors:** Rui Guo, Kaiyao Zhang, Xuze Gao, Xin Jing, Sijia Guo, He Zang, Yuxuan Song, Kunze Li, Peiyuan Zou, Mengjun Chen, Ying Wu, Zhijian Huang, Zhongmin Fu, Dafu Chen

## Abstract

Non-coding RNA (ncRNA) plays an important role in the regulation of immune responses, growth, and development in plants and animals. Here, the identification, characteristic investigation, and molecular verification of circRNAs in *Apis cerana cerana* larval guts were conducted, and the expression pattern of larval circRNAs during *Ascosphaera apis* infection was analyzed. This was followed by exploration of the potential regulatory part of differentially expressed circRNAs (DEcircRNAs) in host immune responses. A total of 3178 circRNAs in the larval guts of *A. c. cerana* were identified, with a length distribution ranging from 15 nt to 96007 nt. Additionally, 45, 33, and 48 up-regulated circRNAs, as well as 110, 62, and 38 down-regulated circRNAs were identified in the *A*.*-apis*-inoculated 4-, 5-, and 6-day-old larval guts in comparison with the corresponding uninoculated larval guts. These DEcircRNAs were predicted to target 29, 25, and 18 parental genes, which were relative to 12, 20, and 17 GO terms as well as 144, 114, and 61 KEGG pathways, including five cellular and four humoral immune pathways containing melanization, phagosomes, lysosomes, endocytosis, apoptosis, MAPK, Ras, and Jak-STAT signaling pathways. Furthermore, complex competing endogenous RNA (ceRNA) regulatory networks were detected as being formed among DEcircRNAs, DEmiRNAs, and DEmRNAs. The target DEmRNAs were engaged in 36, 47, and 47 GO terms as well as 331, 332, and 331 pathways, including six cellular and six humoral immune-related pathways. In total, nineteen DEcircRNAs, five DEmiRNAs, and three mRNAs were included in the sub-networks relative to three antioxidant enzymes, including superoxide dismutase (SOD), catalase (CAT), and glutathione S-transferase (GST). Finally, back-splicing sites within 15 circRNAs and the difference in the 15 DEcircRNAs’ expression between uninoculated and *A*.*-apis*-inoculated larval guts were confirmed utilizing molecular methods. These findings not only enrich our understanding of bee host–fungal pathogen interactions, but also lay a foundation for illuminating the mechanism underlying the DEcircRNA-mediated immune defense of *A. c. cerana* larvae against *A. apis* invasion.

## 1. Introduction

As the most important pollinating insect in nature, honey bees play an important role in pollinating wild plants and crops, and play a pivotal role in the maintenance of biodiversity and ecological balance [1]. *Apis cerana cerana* is an endemic species in China. It is a subspecies of *A. cerana* and one of the major bee species used in beekeeping production. *Ascosphaera apis*, a specialized fungal pathogen of honey bee larvae, causes chalkbrood disease, which causes a dramatic decrease in colony strength and productivity [2].

Circular RNAs (circRNAs), a large class of non-coding RNAs that are produced by a downstream splice-donor site that is covalently linked to an upstream splice-acceptor site, lack the 5′ cap and the poly(A) tail [3]. Arnberg et al. [4] first observed yeast mitochondrial circRNA by electron microscopy in 1980. However, due to the limitations of technical means, the research on circRNAs was nearly stagnant for a long time afterward. With the development of high-throughput sequencing technology and the continuous improvement of RNA cyclization prediction algorithms, the systematic identification of circRNAs at the whole transcriptome level has been achieved. In recent years, circRNAs have been shown to play a wide range of roles in gene regulation at the transcriptional level, post-transcriptional level, and translation level. This includes interacting with RNA polymerase II to promote host gene transcription, acting as molecular sponges to regulate miRNA activity and function, influencing variable gene shearing, and encoding small peptides to translate proteins. Abundant circRNAs have been identified and reported in increasing numbers of animals, plants, and microorganisms such as humans [5, 6], mice [7], rice [8], *Arabidopsis thaliana* [9], *Nosema ceranae* [10], and *Magnaporthe oryzae* [11]. However, research on insect circRNAs is currently still in the preliminary stage. There has been limited documentation for a small number of insects including *Culex pipiens pallens* [12], *Apis mellifera* [*13*], *Bombyx mori* [14], *Drosophila* [15], and *Laodelphax striatellus* [16]. Accumulating evidence has demonstrated that circRNAs are engaged in the regulation of interactions between insects and pathogens/parasites [12, 16-18]. For example, based on high-throughput sequencing of the midgut tissues of both uninfected and BmCPV-infected *B. mori*, Hu et al. [17] detected that 400 host circRNAs were significantly differentially expressed following BmCPV infection. In addition, they found that circRNA_9444, circRNA_8115, circRNA_4553, and circRNA_6649 could act as “molecular sponges” to absorb bmo-miR-278-3p, which negatively regulated the insulin-related peptide binding protein 2 gene and thus participated in the BmCPV-*B. mori* interactions. Zhang et al. [16] identified 2523 circRNAs in the RBSDV-infected midgut tissues of *Laodelphax striatellus* by transcriptome sequencing and bioinformatics. They further screened eight up-regulated and five down-regulated circRNAs, indicative of the involvement of these circRNAs in host response to RBSDV infection. Compared with *Drosophila* and *B. mori*, understanding of circRNA in honey bees is fairly limited. Huang et al. [19] identified 33, 144, and 211 DEcircRNAs by resolving the differential expression profiles of *Apis mellifera ligustica* workers at 1, 5 and 10 days after dinotefuran exposure. Further DEcircRNA–miRNA–mRNA regulatory network analysis showed that circ_0008898 and circ_0001829 were potentially involved in the host immune response as “molecular sponges” of miRNAs. Previously, we discovered 10833 and 9589 circRNAs in *Apis mellifera ligustica* and *Apis cerana* workers’ midguts, and investigated the potential roles of circRNAs in modulating the development of the midguts [20, 21]. Our previous studies suggested that circRNAs were crucial regulators in responses of bee host to infections by fungal pathogens such as *N. ceranae* and *A. apis*. For example, Chen et al. [22] predicted 8199 and 8711 circRNAs in the midgut tissues of *A. m. ligustica* workers at 7 days and 10 days post inoculation (dpi) with *N. ceranae*. They found that 16 circRNAs were highly conserved among *Homo sapiens, A. m. ligustica*, and *Apis cerana cerana*, and deciphered the expression pattern and regulatory role of DEcircRNAs in the host response. Ye et al. [13] identified 2083 circRNAs in the *A. m. ligustica* larval guts and analyzed the structural property of circRNAs, followed by investigation of the roles of DEcircRNAs in modulating the host immune response. However, so far, studies on the whether and how circRNAs regulate interactions between *A. cerana* larvae and *A. apis* is completely unknown.

In this current study, on the basis of previously obtained high-quality RNA-seq datasets, transcriptome-wide identification of circRNAs in *A. c. cerana* larval guts were conducted, and the expression profile of circRNAs was analyzed. This was followed by investigation of the potential roles of DEcircRNAs in regulating the host response to *A. apis* infection. DEcircRNAs and corresponding target genes relevant to antioxidant enzymes and immune responses in the host guts were further analyzed, and ultimately, validation of back-splicing sites and expression trends of DEcircRNAs were performed. Our data could not only lay a foundation for clarifying the mechanisms underlying circRNA-mediated responses of *A. c. cerana* larvae to *A. apis* infection, but also offer new insights into interactions between *A. c. cerana* larvae and *A. apis*.

## 2. Materials and Methods

### 2.1 Fungi and bee larvae

*A. apis* was previously isolated from chalkbrood mummies and conserved at the Honey Bee Protection Laboratory of the College of Animal Sciences (College of Bee Science), Fujian Agriculture and Forestry University, Fuzhou, China [23-25]. *A. c. cerana* worker larvae were derived from three colonies reared in the teaching apiary of the College of Animal Sciences (College of Bee Science) at Fujian Agriculture and Forestry University, Fuzhou, China.

### 2.2 RNA-seq data source

In our previous study, the 4-, 5- and 6-day-old larval guts inoculated with *A. apis* spores (AcT1, AcT2, and AcT3 groups) and the uninoculated 4-, 5- and 6-day-old larval guts (AcCK1, AcCK2, and AcCK3 groups) were prepared and subjected to RNA isolation and strand-specific cDNA-library-based RNA-seq [24, 26,27]. Briefly, (1) the total RNA of three samples in each group was extracted with the TaKaRa MiniBEST Universal RNA Extraction Kit (TaKaRa, Japan), and linear RNA was then removed with Rnase R (GENESEED, China) after removal of rRNA. (2) The obtained circRNA fragments were fragmented into small fragments using a fragmentation buffer, and the first-strand cDNA was synthesized using random hexamer primers and reverse transcription. (3) Second-strand cDNA synthesis was carried out with DNA polymerase I and RNase H, and the double-stranded cDNAs were then purified using the QiaQuick PCR extraction kit (QIAGEN, Germany); the required fragments were then purified by agarose gel electrophoresis followed by enrichment through PCR amplification. (4) The constructed cDNA libraries were sequenced on the Illumina HiSeq™ 2500 platform (GeneDenovo Co., Guangzhou, China. Following our previously described method [21, 28], the produced raw data were subjected to quality control to gain high-quality clean reads, which were used for downstream bioinformatic analyses. Clean reads were mapped to the *A. cerana* reference genome (assembly ACSNU-2.0), followed by identification of circRNAs according to the described method by Guo et al. [29]. Different types of circRNAs were then calculated and raw data were deposited in the NCBI SRA database under the BioProject number: PRJNA562787.

### 2.3 Screening of DEcircRNAs

The expression level of each circRNA was calculated by RPM (reverse splicing node reads per million mapping) method. Following the standard of *P*<0.05 and fold change (FC)≥ 2, DEcircRNAs in every comparison groups were screened. Venn analysis of DEcircRNAs was then performed using the OmicShare platform (https://www.omicshare.com/).

### 2.4 Prediction and annotation of parental genes of DEcircRNAs

Following the method reported by Chen et al. [21], the parental genes of DEcircRNAs were predicted by mapping the anchor reads at both ends of DEcircRNAs to the *A. cerana* reference genome (assembly ACSNU-2.0) using Bowtie 2 software [30] with default parameters. If both ends of one circRNA were aligned to the same gene, this gene was regarded as the source gene of the circRNA. Next, the parental genes were annotated to GO (http://www.geneontology.org/) and KEGG (https://www.kegg.jp/) databases by the BLAST tool with default parameters.

### 2.5 Source of small RNA-seq datasets

In another previous study, *A*.*-apis*-inoculated and uninoculated 4-, 5-, and 6-day-old larval guts of *A. c. cerana* were prepared, followed by RNA isolation, cDNA library construction, and sRNA-seq. Quality control of the raw data was then carried out to gain high-quality clean tags [31], which were used for target prediction in this study. The raw data are available in the NCBI SRA database under the BioProject number: PRJNA562787.

### 2.6 Analysis of the ceRNA regulatory network

By using Targetfinder [32] and mirTarBase software, the potential targeting relationships between DEcircRNA and DEmiRNAs, as well as between DEmiRNAs and DEmRNAs, were predicted. Following the predicted targeting relationships, DEcircRNA–DEmiRNA–DEmRNA regulatory networks were constructed and then visualized by Cytoscape v.3.2.1 software [33] with default parameters. Next, the targets were mapped to the Nr, GO, and KEGG databases with the BLAST tool.

### 2.7 Investigation of antioxidant enzyme-associated DEcircRNAs and the corresponding regulatory network

Based on the Nr annotations in Section 2.5, similarly, the potential targeting relationships between antioxidant-enzyme-associated mRNAs and DEmiRNAs, as well as between DEmiRNAs and DEcircRNAs, were predicted with Targetfinder [32] and mirTarBase software. Further, DEcircRNA–DEmiRNA–mRNA regulatory networks were constructed and then visualized by Cytoscape v.3.2.1 software [33].

### 2.8 Investigation of immune response-related DEcircRNAs and the corresponding regulatory network

The potential targeting relationships between immune-defense-related DEmRNAs and DEmiRNAs, as well as DEmiRNAs and DEcircRNAs, were predicted with Targetfinder [32] and mirTarBase software. On the basis of the predicted targeting relationships, DEcircRNA– DEmiRNA–DEmRNA regulatory networks were constructed and then visualized by Cytoscape v.3.2.1 software [33].

### 2.9 Prediction and analysis of DEcRNAs with coding potential

The IRES contained in DEcircRNA were predicted with IRESfinder software [34]. And the open-reading frames (ORFs) were predicted using ORFfinder [35]. The ORFs were annotated to the GO and KEGG databases.

### 2.10 PCR amplification and Sanger sequencing of circRNAs

To confirm the authenticity of circRNAs, 15 circRNAs were randomly selected for PCR amplification and Sanger sequencing, including novel_circ_000983, novel_circ_001484, novel_circ_002377, novel_circ_002038, novel_circ_002313, novel_circ_002486, novel_circ_000504, novel_circ_001175, novel_circ_002439, novel_circ_000526, novel_circ_000446, novel_circ_001799, novel_circ_001391, novel_circ_002045, and novel_circ_002378. Across the back-splicing sites, a divergent primer was designed using DNAMAN software (shown in Table S1) and synthesized by Sangon Biotech (Shanghai) Co., Ltd. The TaKaRa MiniBEST Universal RNA Extraction Kit (TaKaRa, Japan) was used to extract the total RNA from the total RNA of larval gut samples in the AcCK1, AcCK2, AcCK3, AcT1, AcT2, and AcT3 groups. This was followed by digestion of linear RNA with 3 U/mg RNase R to enrich circRNAs. The template was treated at 37°C for 15 min, and the cDNA of circRNA was obtained by reverse transcription with random primers. These were then used as templates for PCR amplification, which was conducted on a T100 thermal cycler (BioRad, USA). The PCR reaction system consisted of 10 μL of PCR Mix, 2 μL of the DNA template, 1 μL each of upstream and downstream primers (2.5 pmol/μL), and 6 μL of sterile water. The reaction procedure was set as: 95°C for 5 min, 95°C for 30 s, and 60°C for 30 s, for 34 cycles; then, 72°C for 2 min. The amplified products were detected by 1.5% agarose gel electrophoresis with Ultra GelRed staining (Vazyme, China). This was followed by purification of the target fragments with the FastPure Gel DNA Extraction Mini Kit (Vazyme, China) and then Sanger sequencing by Sangon Biotech (Shanghai) Co., Ltd.

### 2.11 RT-qPCR detection of DEcircRNAs

To further verify the reliability of circRNA sequencing data, five circRNAs were randomly selected from each of the three comparison groups for RT-qPCR validation. The AcCK1 vs. AcT1 group included novel_circ_000983, novel_circ_001484, novel_circ_000882, novel_circ_002486, and novel_circ_002377; The AcCK2 vs. AcT2 group included novel_circ_000504, novel_circ_000405, novel_circ_000526, novel_circ_001175, and novel_circ_002439; The AcCK3 vs. AcT3 group included novel_circ_002378, novel_circ_000102, novel_circ_001799, novel_circ_002486, and novel_circ_001391. The total RNA obtained was divided into two portions: one portion was digested with RNase R to enrich circRNA, and the resulting cDNA obtained by reverse transcription with random primers was used as the templates for RT-qPCR detection of DEcircRNAs; the other portion was subjected to reverse transcription with Oligo dT primers, and the resulting cDNA were used as the templates for RT-qPCR detection of the internal refence gene actin (Gene ID: XM_017059068.2). There were three parallel samples and each experiment was repeated three times. The reaction system followed the method of Ye et al [13]. The reaction was conducted on Applied Biosystems® QuantStudio 3 (ABI, USA) following the conditions: 95°C pre-denaturation for 5 min, 95°C denaturation for 15 s, and 60°C annealing and extension for 30 s, with a total of 40 cycles of qPCR reaction. The relative expression level of each DEcircRNA was calculated using the 2^-∆∆Ct^ method. Data were shown as the mean ± standard deviation (SD) and subjected to Student’s *t*-test by Graph Prism 8 software (“ns” represents *P*>0.05, “*” represents *P*<0.05; “**” represents *P*<0.01; “***” represents *P*<0.001). Details of RT-qPCR primers are presented in Table S1.

## 3. Results

### 3.1. Quality control of deep sequencing data

In total, 73 830 148, 96 586 212, 94 552 744, 76 672 564, 90 954 858, and 83 418 832 raw reads were generated in the AcCK1, AcCK2, AcCK3, AcT1, AcT2, and AcT3 groups, respectively (Table S2). After undergoing strict quality control, 73 775 592, 96 513 798, 94 495 000, 76 593 924, 90 870 608, and 83 339 288 clean reads were identified, with Q30 above 93.42% (Table S2). The results indicated that the sequencing data were of high quality.

### 3.2 Identification and characterization of the A. c. cerana circRNAs

Based on the total clean reads, 3178 *A. c. cerana* circRNAs were discovered (Table S3), with a length distribution ranging from 15 nt to 96 007 nt; those circRNAs distributed among 301∼400 nt represented the largest group (757, 23.81%)(Figure 1A). The identified circRNAs included five types, among which exonic circRNA (66.76%) was the most abundant type, followed by exon– intron circRNA (15.73%), antisense circRNA (12.30%), intronic circRNA (2.64%), and intergenic region circRNA (2.55%). Additionally, it was found that NW_016019774.1 was the most widely distributed chromosome by circRNAs in the AcCK1 group, while NW_016017967.1 was the most enriched by circRNAs in the AcCK2 and AcCK3 groups; NW_016017455.1 was the most commonly distributed chromosome by circRNAs in the AcT1 group, whereas NW_016017967.1 was the most enriched by circRNAs in the AcT2 and AcT3 groups (Figure 1C).

**Figure 1.**
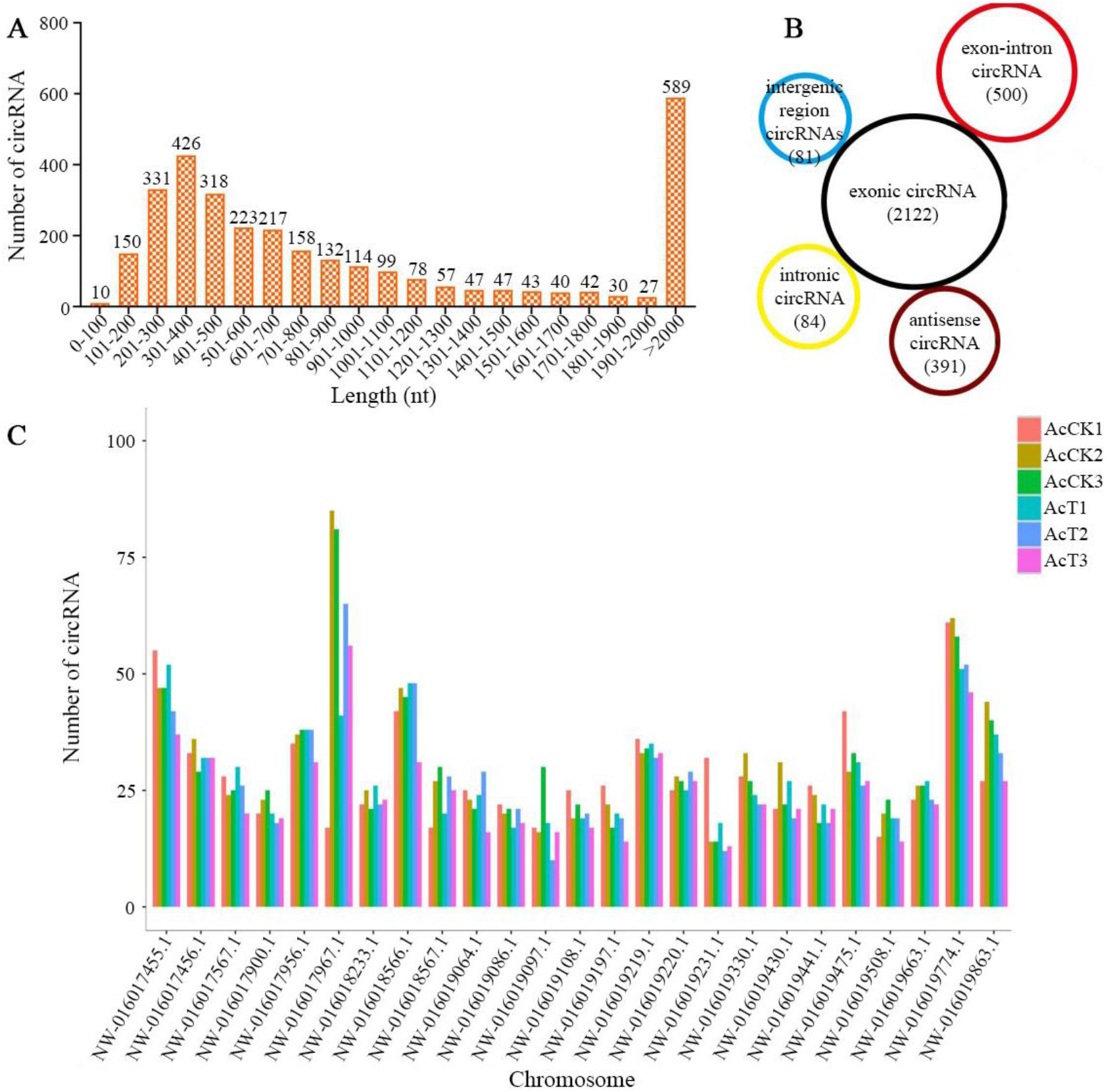
Characteristics of circRNAs identified in *A. c. cerana*. (**A**) Length distribution; (**B**) types, the numbers of various types of circRNAs are presented in brackets; (**C**) chromosome distribution.

### 3.3 Expression profile of circRNAs engaged in larval response to A. apis infection

In the 4-day-old comparison group, 155 DEcircRNAs were screened, including 45 up-regulated and 110 down-regulated circRNAs; in the 5- and 6-day-old comparison groups, 33 and 48 up-regulated circRNAs, as well as 62 and 38 down-regulated circRNAs were detected, respectively (Figure 2A). Venn analysis showed that four up-regulated and one down-regulated circRNAs were shared by the above-mentioned three comparison groups (Table 1), whereas the numbers of unique circRNAs were 125, 70, and 66, respectively (Figure 2B, Table S4).

**Table 1.** Detailed information about circRNAs shared by 4-, 5-, and 6-day-old comparison groups.

| ID | Source gene | Chromosome | Strand | Genomic start | Genomic end | Spliced length | Annotated type |
| --- | --- | --- | --- | --- | --- | --- | --- |
| novel_circ_001502 | ncbi_107996165 | NW_016019064.1 | - | 1633180 | 1635297 | 2118 | antisense |
| novel_circ_002548 | ncbi_108001011 | NW_016019508.1 | + | 95288 | 95775 | 488 | one_exon |
| novel_circ_002606 | ncbi_108001152 | NW_016019530.1 | - | 119107 | 119379 | 273 | one_exon |
| novel_circ_002608 | ncbi_108001152 | NW_016019530.1 | - | 119108 | 119311 | 204 | one_exon |
| novel_circ_002609 | ncbi_108001152 | NW_016019530.1 | - | 119108 | 119380 | 273 | one_exon |

**Figure 2.**
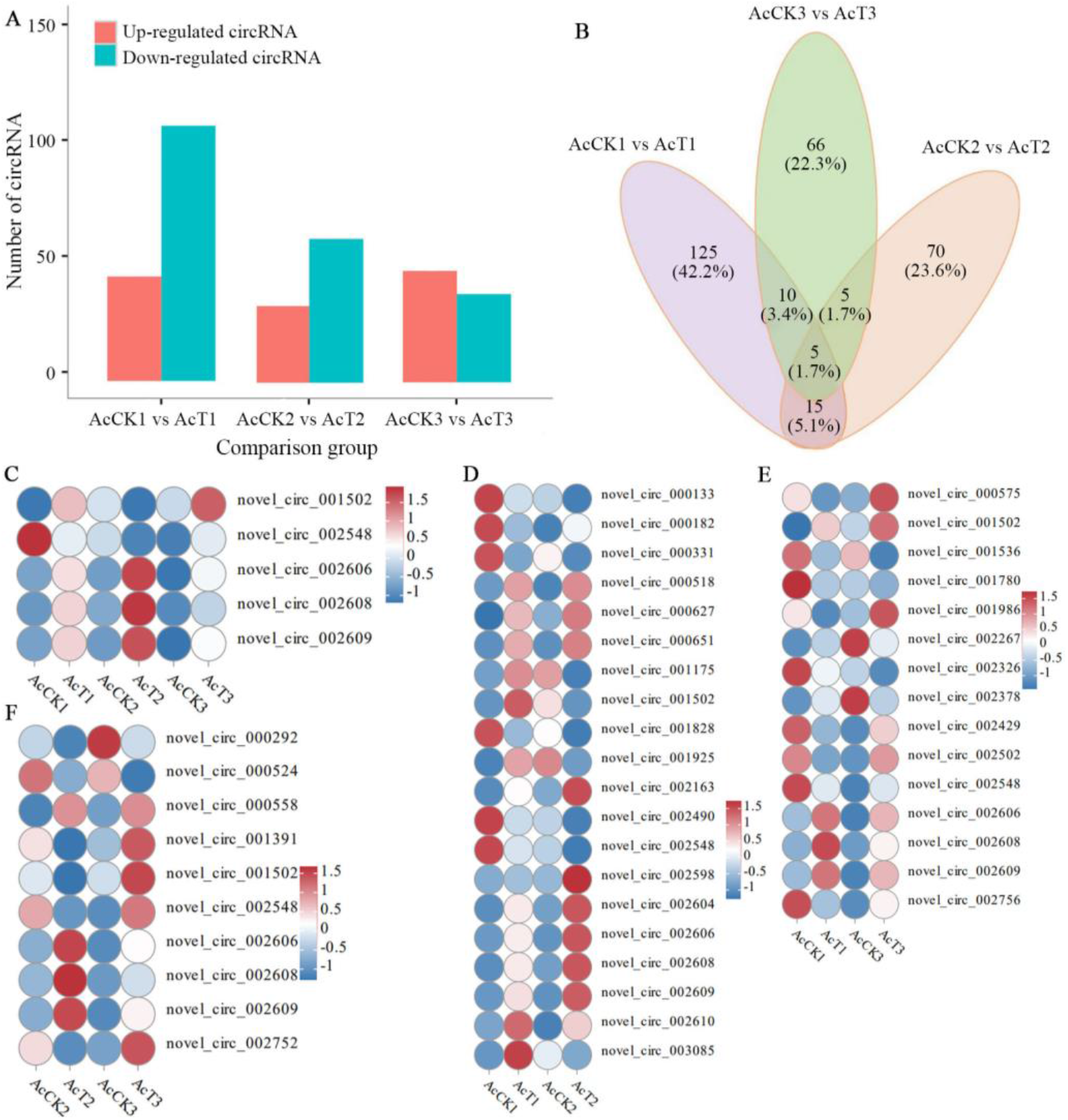
Expression clustering of DEcircRNAs in three comparison groups. (**A**) Number statistics of DEcircRNAs in three comparison groups; (**B**) Venn analysis of DEcircRNAs in three comparison groups; (**C**) Expression clustering of shared DEcircRNAs by three comparison groups; (**D**) Expression clustering of shared DEcircRNAs by AcCK1 vs. AcT1 and AcCK2 vs. AcT2 comparison groups; (**E**) Expression clustering of shared DEcircRNAs by AcCK1 vs. AcT1 and AcCK3 vs. AcT3 comparison groups; (**F**) Expression clustering of shared DEcircRNAs by AcCK2 vs. AcT2 and AcCK3 vs. AcT3 comparison groups.

### 3.4 Annotation of the DEcircRNAs’ parental genes

It’s predicted that 29 parental genes of DEcircRNAs in the 4-day-old comparison group were annotated to 12 biological process-associated GO terms such as cellular process and metabolic process, eight cellular component-associated terms such as membrane part and cell, and six molecular function-associated terms such as catalytic activity and transporter activity (Figure 3A); the parental genes were also enriched in 144 KEGG pathways, such as lysosome, melanogenesis, and PI3K-Akt signaling pathway (Figure 3B). Comparatively, twenty-five parental genes of DEcircRNAs in the 5-day-old comparison group were engaged in 20 terms (cellular process, binding, and single-organism process etc.) (Figure 3C) and 114 pathways (phagosome, apoptosis-fly, MAPK signaling pathway-fly etc.) (Figure 3D). In the 6-day-old comparison group, 18 parental genes were involved in 17 terms (binding, catalytic activity, cell etc.) (Figure 3E) and 61 pathways (apoptosis, endocytosis, Jak-STAT signaling pathway etc.) (Figure 3F). The numbers of parental genes relevant to cellular and humoral immune pathways were summarized in Figure 3G. Intriguingly, MAPK signaling pathway was observed to be enriched by parental genes of DEcircRNAs in the aforementioned three comparison groups.

**Figure 3.**
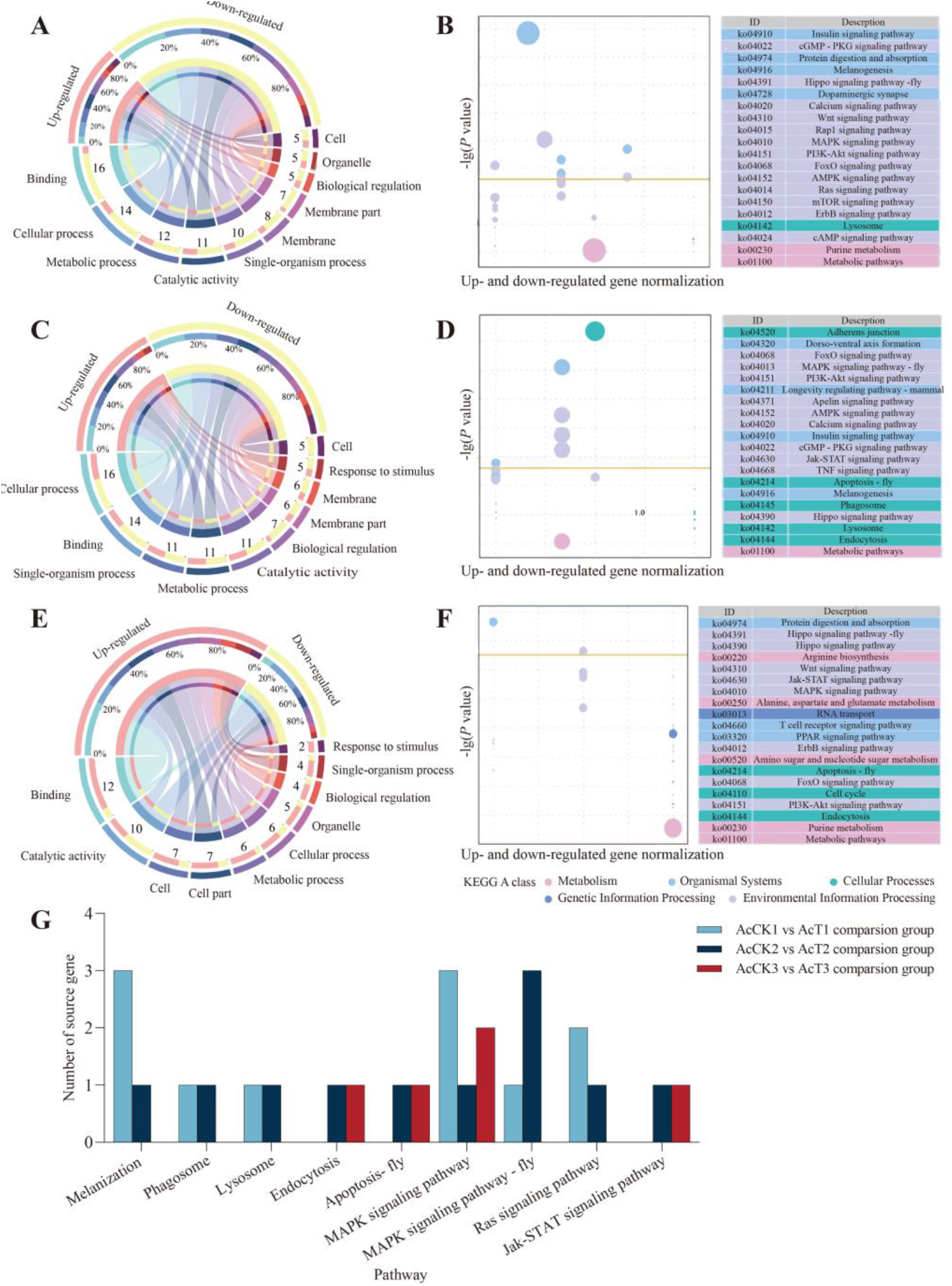
Annotation of parental genes of DEcircRNAs in three comparison groups. (**A, C, E**) Loop graphs of GO terms annotated by parental genes; (**B, D, F**) KEGG pathways annotated by parental genes; (**G**) Number statistics of parental genes relative to cellular and humoral immune pathways.

### 3.5 Analysis of ceRNA regulatory networks

CeRNA regulatory network analysis demonstrated that 41, 31, and 59 DEcircRNAs in the above-mentioned three comparison groups could target nine, 26, and 54 DEmiRNAs (Figure 4), further targeting 760, 4464, and 5015 DEmRNAs, respectively. Additionally, a subseries of DEcircRNAs can simultaneously target multiple DEmiRNAs, e.g., novel_circ_002084, novel_circ_002977, and novel_circ_001648 in the 4-day-old comparison group could target four, three, and three DEmiRNAs. Meanwhile, some DEmiRNAs could also be targeted by several DEcircRNAs at the same time, e.g., miR-1277-x, novel-m0006-5p, and miR-1344-x could be targeted by fifteen, ten, and night DEcircRNAs in the 6-day-old comparison groups.

**Figure 4.**
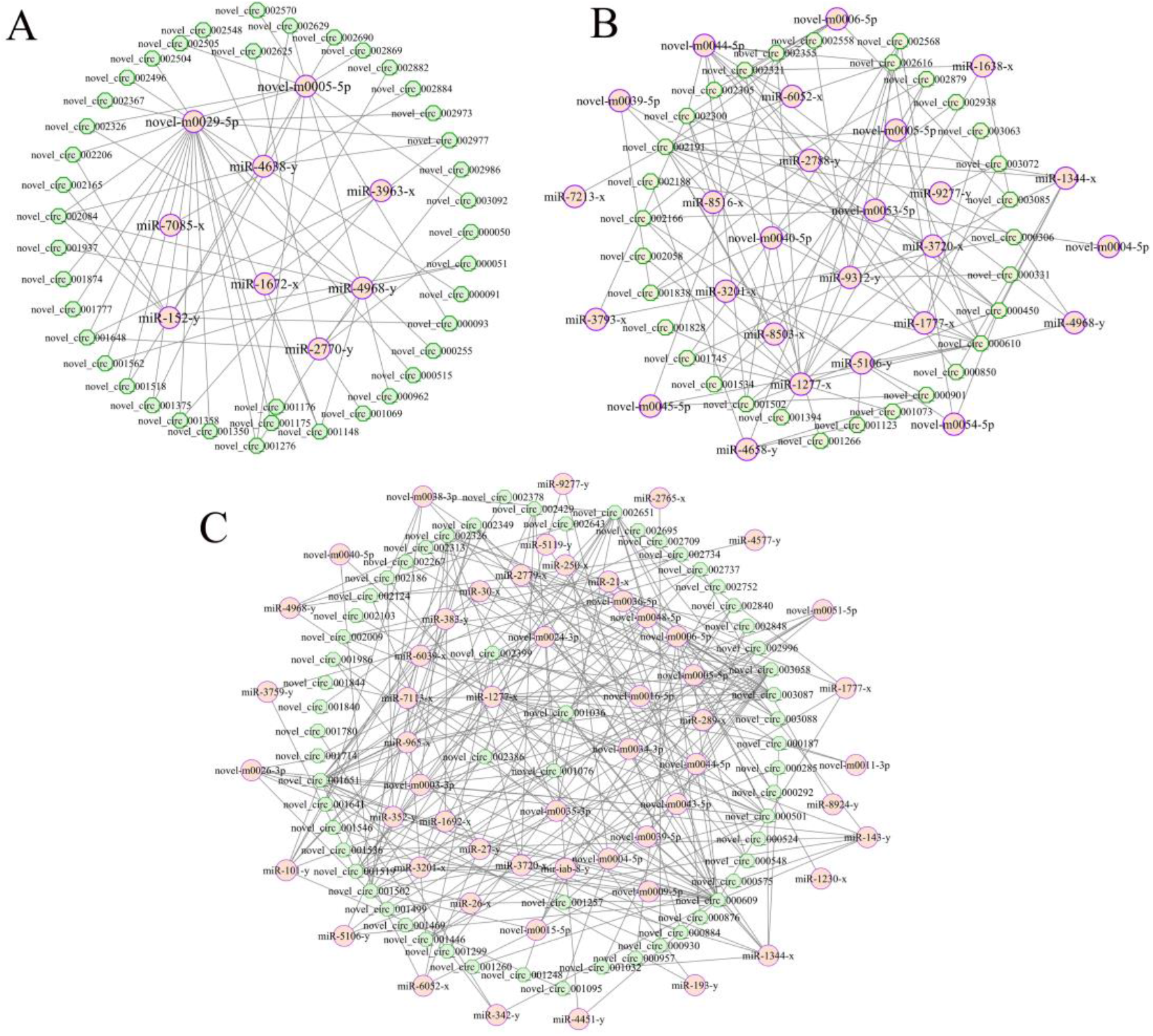
Regulatory networks between DEcircRNAs and DEmiRNAs. (**A–C**) DEcircRNA–DEmiRNA networks in AcCK1 vs. AcT1, AcCK2 vs. AcT2, and AcCK3 vs. AcT3 comparison groups. Hexagons represent DEcircRNAs, while circles represent DEmiRNAs.

Targets in the 4-day-old comparison group were involved in 36 GO terms (cellular process, cell, binding, etc.) (Figure 5A) and 331 KEGG pathways (metabolic pathways, endocytosis, RNA transport, etc.) (Figure 6A). In contrast, targets in the 5-day-old comparison group were engaged in 47 terms (cellular process, cell, binding, etc.) (Figure 5B) and 332 pathways (metabolic pathways, endocytosis, MAPK signaling pathway, etc.) (Figure 6B). In the 6-day-old comparison group, these targets were relevant to 47 terms (metabolic process, cell, binding, etc.) (Figure 5C), as well as 331 pathways (metabolic pathways, RNA transport, Wnt signaling pathway, etc.) (Figure 6C).

**Figure 5.**
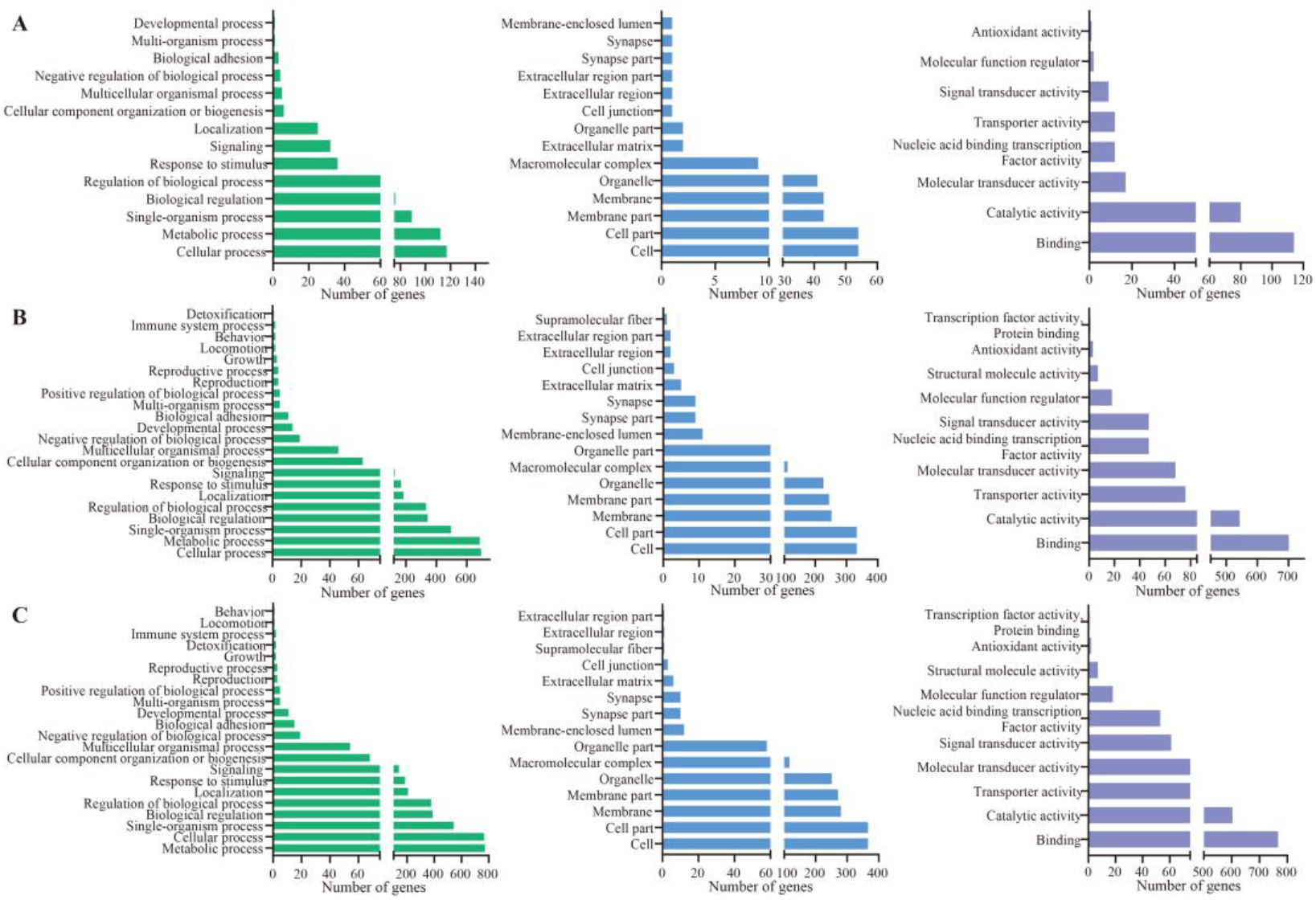
GO terms annotated by targets in DEcircRNA-involved ceRNA regulatory networks. (**A–C**) Biological process, cell component, and molecular-function-related terms annotated by targets in the AcCK1 vs. AcT1, AcCK2 vs. AcT2, and AcCK3 vs. AcT3 comparison groups.

**Figure 6.**
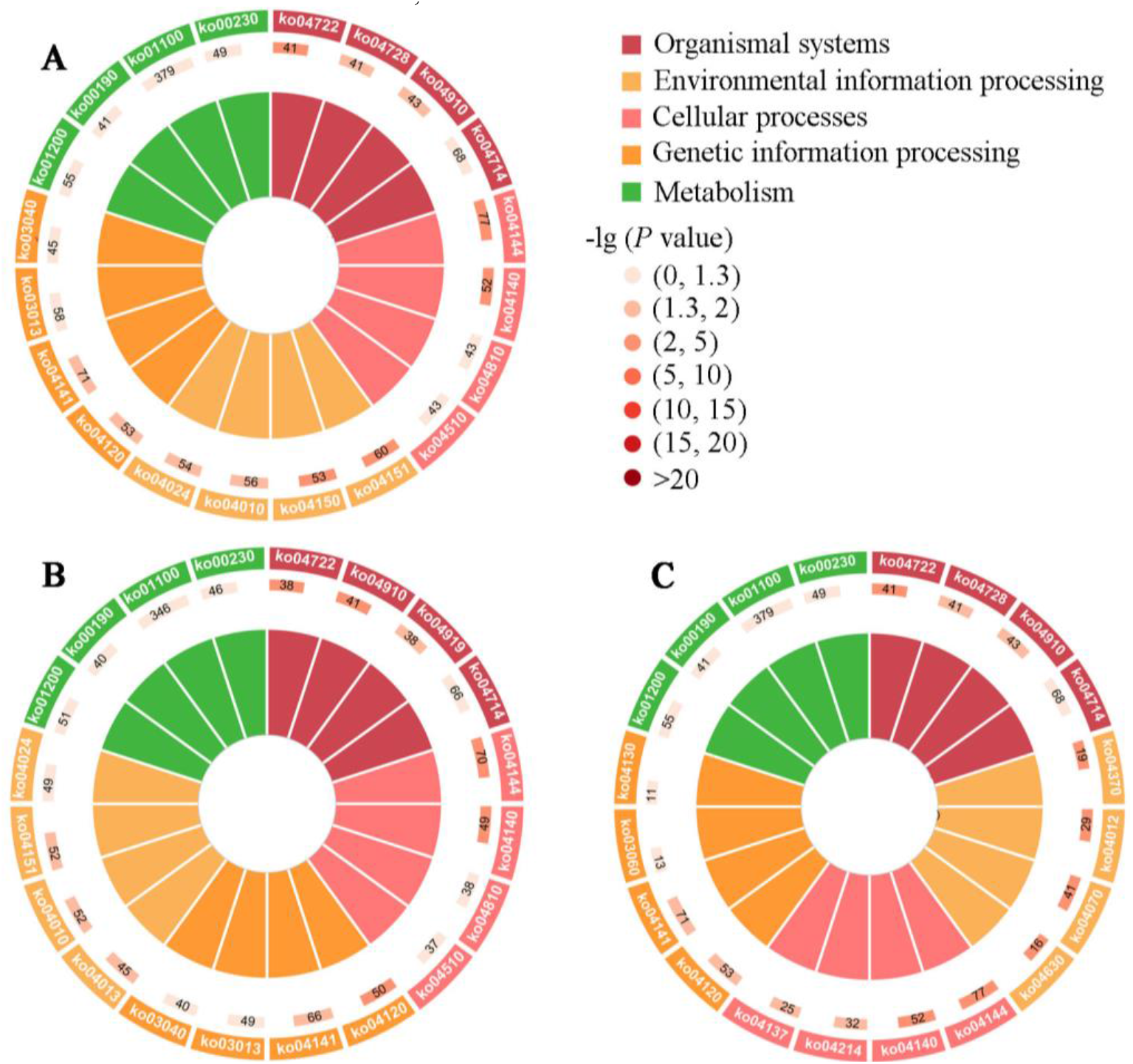
KEGG pathways enriched by targets in DEcircRNA-involved ceRNA regulatory networks. (**A–C**) Pathways enriched by targets in the 4-, 5-, and 6-day-old comparison groups.

### 3.6 Investigation of Antioxidant-Enzyme-Associated DEcircRNAs and the corresponding regulatory network

Further analysis demonstrated that nineteen DEcircRNAs, five DEmiRNAs, and three mRNAs shared by the 4-, 5-, and 6-day-old comparison groups were included in the sub-networks relative to three antioxidant enzymes including superoxide dismutase (SOD), catalase (CAT), and glutathione S-transferase (GST) (Figure 7, see also Table S5). In detail, three DEcircRNAs potentially targeted one DEmiRNA, further targeting one mRNA associated with superoxide dismutase (Figure 7, see also Table S5); ten DEcircRNAs putatively targeted two DEmiRNAs, further targeting one mRNA related to catalase (Figure 7, see also Table S5); six DEcircRNAs potentially targeted two DEmiRNAs, further targeting one mRNA relevant to glutathione S-transferase (Figure 7, see also Table S5).

**Figure 7.**
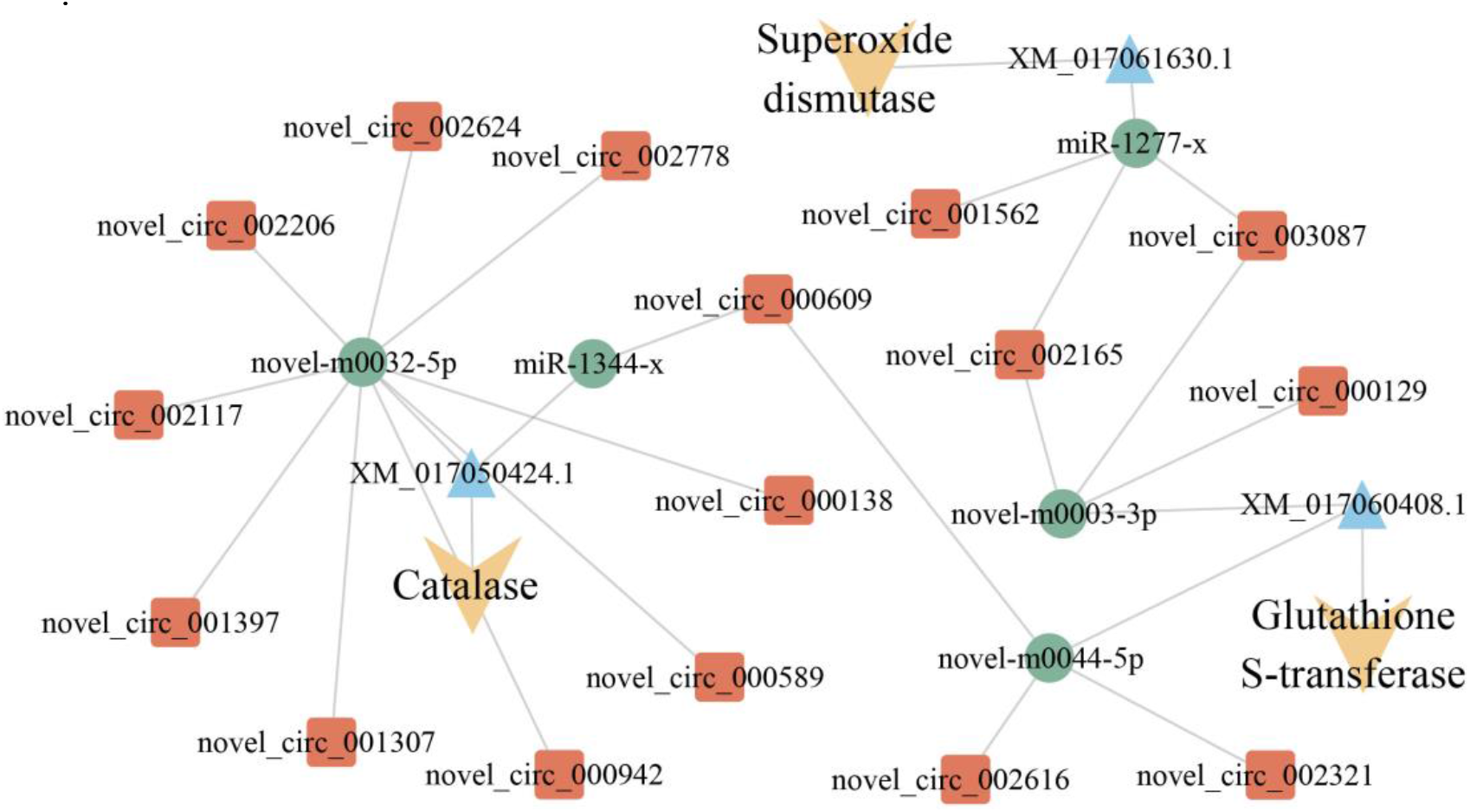
CeRNA regulatory networks of antioxidant-enzyme-associated DEcircRNAs.

### 3.7 Analysis of immune-defense-related DEcircRNAs and the corresponding regulatory network

It was observed that immune-defense-related sub-networks included 56 DEcircRNAs, 13 DEmiRNAs, and 49 DEmRNAs for the 4-, 5-, and 6-day-old comparison groups, respectively (Figure 8, see also Table S6). In detail, 51 DEcircRNAs putatively targeted 9 DEmiRNAs, further targeting 31 DEmRNAs involved in 6 cellular immune-related pathways, including apoptosis, melanogenesis, endocytosis, autophagy-animal, apoptosis-fly, and insect hormone biosynthesis (Figure 8, see also Table S6); 51 DEcircRNAs potentially targeted 10 DEmiRNAs, further targeting 22 DEmRNAs engaged in 6 humoral immune-related pathways, such as the Toll and Imd, Toll-like receptors, NF-kappa B, Jak-STAT, and MAPK signaling pathways (Figure 8, see also Table S6). In addition, 46 circRNAs were found to be involved in the regulatory networks regarding both cellular and humoral immune responses.

**Figure 8.**
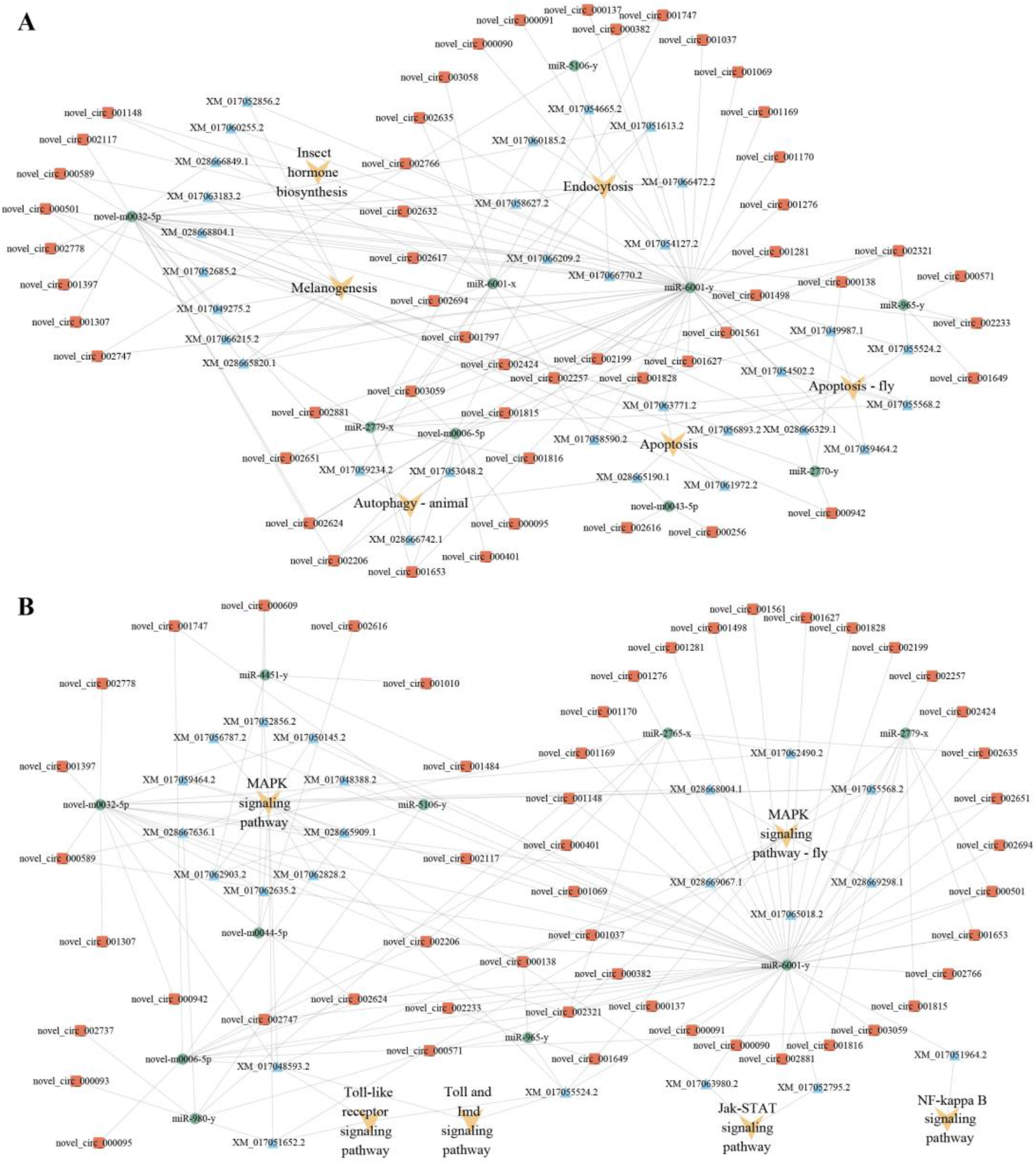
CeRNA regulatory networks of cellular and humoral immune-related DEcircRNAs.

### 3.8 Investigation of the protein-coding ability of DEcircRNA

In the 4-day-old comparison group, 27 IRESs and 61 ORFs within DEcircRNAs were discovered; these ORFs were engaged in 23 GO terms such as cellular processes and signaling (Table S7), as well as 107 KEGG pathways such as the insulin signaling pathway and melanogenesis (Table S8). In the 5-day-old comparison group, 26 IRESs and 52 ORFs within DEcircRNAs were identified; these ORFs were relative to 20 functional terms including binding and the response to stimulus (Table S7), as well as 71 pathways including metabolic pathways and the MAPK signaling pathway (Table S8). Additionally, 24 IRESs and 40 ORFs within DEcircRNAs in the 6-day-old comparison group were detected; these ORFs were involved in 17 functional terms such as catalytic activity and cell processes (Table S7), as well as 16 pathways such as endocytosis and apoptosis (Table S8).

### 3.9 Molecular verification of Back-Splicing Sites within DEcircRNAs

Five randomly selected DEcircRNAs from each comparison group were subjected to PCR amplification followed by Sanger sequencing. The results confirmed the authenticity of the back-splicing sites in these DEcircRNAs (Figure 9).

**Figure 9.**
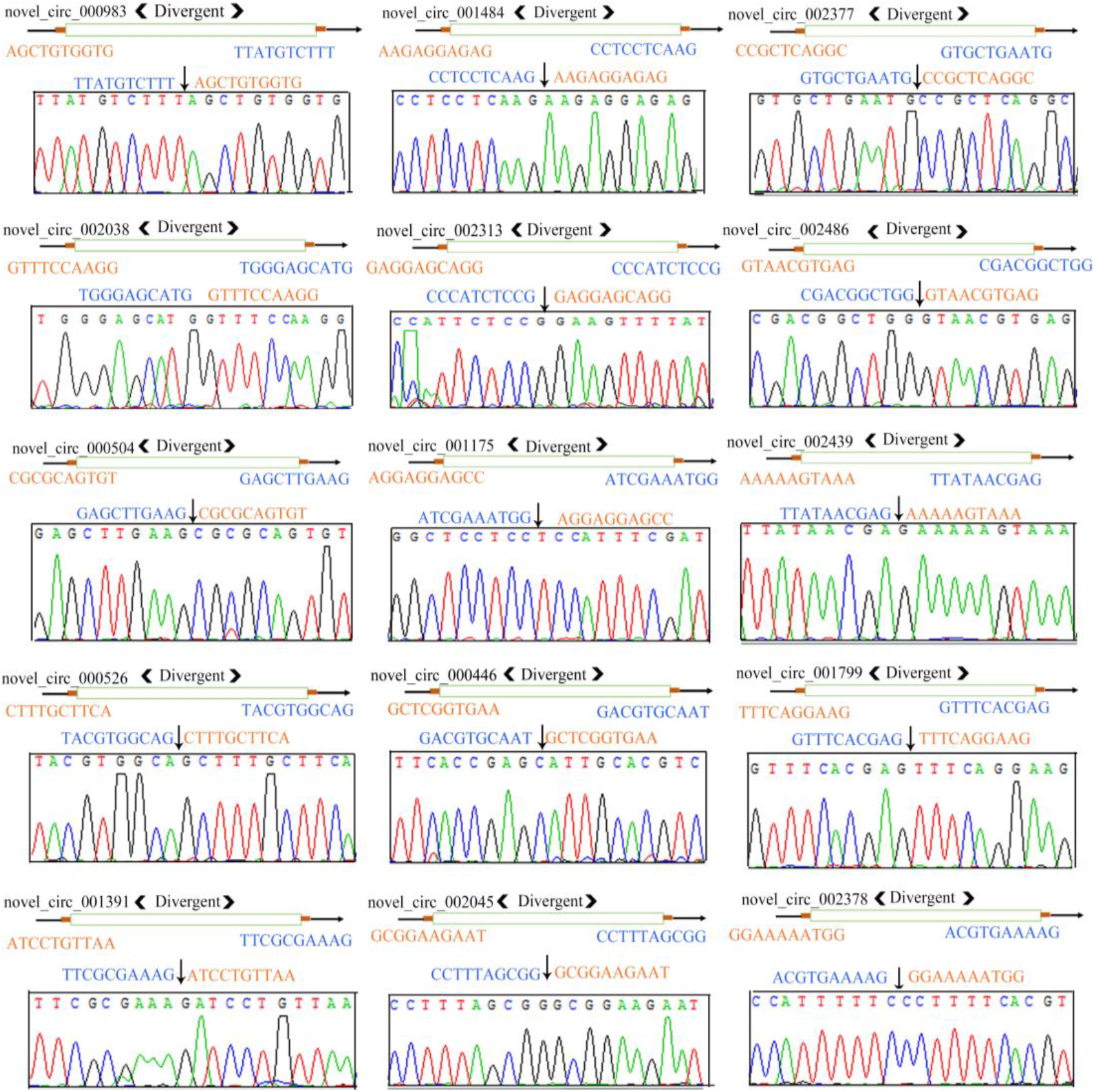
Sanger sequencing of amplification products from 15 circRNAs. “<” and “>” indicate the direction of amplification using divergent primers; “→” indicates the transcriptional directions of circRNAs; “→” indicates the back-splicing sites within circRNAs.

### 3.10 RT-qPCR detection of DEcircRNAs

Further, RT-qPCR of the aforementioned 15 DEcircRNAs were conducted. The results indicated that the expression trends between *A*.*-apis*-inoculated and uninoculated 4-, 5-, and 6-day-old larval guts were in accordance with those in the prediction results, based on deep sequencing data (Figure 10), thus validating the reliability of the transcriptome datasets used in this work.

**Figure 10.**
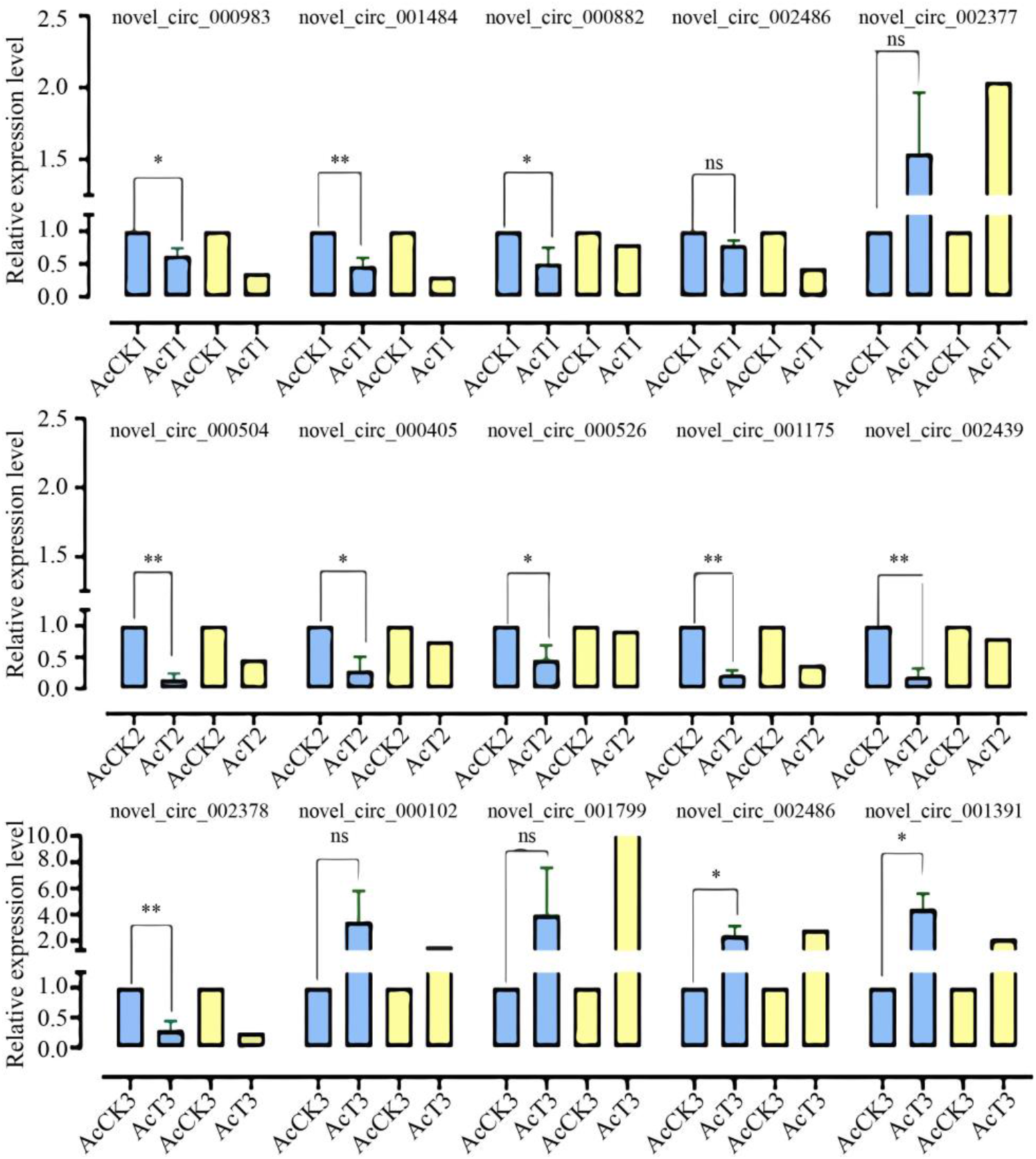
RT-qPCR detection of 15 DEcircRNAs. The experimental data were presented as mean ± SD and subjected to Student’s t test; ns: *P*>0.05; *: *P*<0.05; **: *P*<0.01.

## 4 Discussion

Our group previously identified 3178 circRNAs of infected 4-6-day-old larvae and uninfected 4-6-day-old larvae of *A. c. cerana*. In view of the limited reports on *A. cerana* circRNAs, the circRNAs discovered in this study could further enrich the reservoir of circRNA data for *A. cerana* and provide a valuable resource for future research on circRNAs in *A. c. cerana* and other subspecies belonging to *A. cerana*. The number of circRNAs identified by previous studies in humans [36], mice [36], nematodes [36] and soybean [37] species differed significantly. The number of circRNAs was also significantly different from that of *A. c. cerana* worker larvae identified in this study, indicating that genes of different species can be transcribed to form different amounts of circRNAs. However, differences in library construction methods, bioinformatics algorithms, analysis software setup parameters, individuals of different species, and sequencing tissues in different studies may also be important reasons for the significant differences in the identified number of circRNAs. In addition, we found that the back-splicing sites of *A. c. cerana* circRNA contained the conserved splicing signal GT/AG. This was similar to splicing signals discovered in other animals such as humans [38], mice [39], silkworms [17], and *Chiloscyllium plagiosum* [40], indicating that this splicing signal was conserved among animals. In addition, for circRNAs identified in both *A. apis*-infected and uninfected groups, the major splicing signal was GT/AG, which suggested that the *A. apis* infection could not alter this stable splicing signal. Here, it was observed that circRNAs distributed among 301∼400 nt represented the largest group, analogous to those identified in humans [41], mice [41], and rice [42]. Also, we detected that annotated exonic circRNA was the most abundant type, as shown in Figure 2B. This is similar to the findings from humans [43], *A. m. ligustica* [44], *A. thaliana* [45], and *Zea mays* [46].

In the present study, 155, 95, and 86 DEcircRNAs were discovered in the 4-, 5-, and 6-day-ld comparison groups, respectively (Figure 2), indicating that *A. apis* infection gave rise to the alteration of the overall expression pattern of circRNAs in *A. c. cerana* worker larval guts. These DEcircRNAs were potential regulators in the larval *A. apis* response. Additionally, five DEcircRNAs covering two types of circulation, such as novel_circ_001502, novel_circ_002548, novel_circ_002606, novel_circ_002608, and novel_circ_002609 were shared by the 4-, 5-, and 6-day-old comparison groups. It is inferred that these five shared DEcircRNAs played critical roles in the host response to *A. apis* infection, and thus deserves further investigation.

Accumulating evidence has showed that some circRNAs modulate the transcription of parental genes and further affect an array of biological processes such as immune response and development [47, 48]. In this study, we found that 3178 circRNAs were generated from 1356 parental genes; among these, some parental genes produced only one circRNA, while others produced two or more circRNAs, similar to the reported findings from studies on *Drosophila* [49] and silkworms [14]. In addition, 296 DEcircRNAs were detected as being derived from 225 parental genes. This could mean that a single linear mRNA principally generated only one circRNA, and some could yield two or more circRNAs, suggesting that a complicated circularization mechanism indeed occurred in circRNA biogenesis during *A. apis* infection of *A. c. cerana*. This was annotated to a series of functional terms such as cellular process, metabolic process, cell, organelle, catalytic activity, and binding. In addition, the parental genes mentioned above were also relevant to 184 pathways, including melanogenesis, lysosomes, metabolic pathways, FoxO, hippo, and the MAPK signaling pathway. Together, the results suggest that these DEcircRNAs may have participated in the modulation of the aforementioned life activities by regulating the transcription of corresponding parental genes.

The insulin signaling pathway is evolutionarily conserved and interacts with insect’s innate immunity, while acting synergistically with insect ecdysteroid to induce cellular autophagy and apoptosis in larval tissues. This plays a key role in insect growth and development, reproductive metabolism, and resilience [50]. In insects such as *B. mori* [51], *Drosophila melanogaster* [52], *Schistocerca gregaria* [53], and *Anopheles gambiae* [54], several related insulin-like genes have been identified and shown to be closely associated with growth, development, metabolism, and immune defense. After mutating the gene encoding the insulin receptor in *Drosophila*, Drummond et al. [55] observed that ovarian follicles regressed in *Drosophila* and blocked entry into vitellogenesis. Here, we found six and two parental genes in the 4- and 5-day-old comparison groups, corresponding to seven and three DEcircRNAs, respectively, that could be annotated to the insulin signaling pathway; however, no parental gene in the 6-day-old comparison group was observed as being annotated to this pathway. The insulin receptor catalyzes the autophosphorylation and activation of the intermediary molecule growth-factor-receptor-linked protein Grb2 and the SOS molecule with guanylate exchange factor activity. This is followed by the activation of the Raf protein, which further activates the MAPK signaling pathway due to its serine/threonine protein kinase activity [56]. In eukaryotes, the MAPK signaling pathway, a vital pathway response to oxidative stress, transfers extracellular information to the interior of the cell, further regulating cell activities in response to external stimuli [57]. As an important member of the MAPK signaling pathway, MKK is involved in modulating cell growth and the immune response [58]. Wang et al. [59] previously identified an *AccMKK6* gene in *A. c. cerana*, and detected that the activities of SOD and POD were significantly reduced and the antioxidant capacity of bees was decreased after knockdown of *AccMKK6* in adult workers via RNAi. Here, three, one, and two parental genes, corresponding to three, one, and two DEcircRNAs, respectively, could be annotated to the MAPK signaling pathway (Figure 3G). Collectively, the results demonstrated that these DEcircRNAs were likely to participate in the host response to *A. apis* infection by regulating the transcription of parental genes relative to the three above-mentioned signaling pathways.

Increasingly, studies have shown that circRNAs are capable of acting as a “molecular sponge” for miRNA to regulate the expression of genes and further affect biological processes such as metabolism and immunity [60-63]. In this current study, 41, 31, and 59 DEcircRNAs in the 4-, 5-, and 6-day-old comparison groups were found to target nine, 26, and 54 DEmiRNAs, respectively (Figure 4). This is suggestive of the potential of these host DEcircRNAs to act as a “molecular sponge” for miRNAs during the infection process of *A. apis*. We observed the significant down-regulation of miR-1277-x in the *A. c. cerana* 6-day-old worker larval gut following *A. apis* infection, and found that miR-1277-x putatively targeted a gene belonging to the JAK/STAT signaling pathway. Here, three up-regulated circRNAs (novel_circ_003058, novel_circ_003088, and novel_circ_002651) were found to simultaneously target miR-1277-x, further targeting a downstream gene encoding the *transcriptional regulator Myc-B*. This implied that the *A. apis* infection resulted in the activation of novel_circ_003058, novel_circ_003088, and novel_circ_002651 in the larval guts, which may attenuate the inhibitory effect of miR-1277-x on the expression of *Myc-B*, thereby modulating the Jak-STAT signaling pathway. However, more efforts are needed to verify this speculation.

Insects possess a suite of antioxidant enzymes and antioxidants with small molecular weights, which form a concatenated response to an onslaught of dietary and endogenously produced oxidants. Antioxidant enzymes such as superoxide dismutase, catalase, glutathione transferase, and glutathione reductase have been characterized in insects [64]. Ling and Zhang [65] documented that CAT mRNA was specifically induced in the presence of chlorpyrifos, suggesting that the intensified CAT enzyme activities contributed to enhancing the antioxidant capacity and population growth of *Nilaparvata lugens*. Li et al. [66] identified 31 cytosolic GSTs genes in *Spodoptera litura*, including *SlGSTd1*, a GSTs gene from the delta cluster. They found that silencing *SlGSTd1* significantly increased the cumulative mortality after fenvalerate treatment for 72 h and cyhalothrin treatment for 48, 60 and 72 h. Also, studies have shown that antioxidant enzymes are involved in the response of insects to pathogenic microorganisms [67]. Vivekanandhan et al. [68] showed, in *Spodoptera litura* larvae, that SOD enzyme levels were increased when the *Metarhizium flavoviride* conidia concentrations were increased, and the insect antioxidant enzymes played a significant role in ROS eradication. Various xenobiotics can cause oxidative stress and the production of ROS in insects, Zhang et al. [69] detected that, after *Beauveria bassiana* infected the larvae of *Spodoptera frugiperda*, the activities of SOD and CAT enzymes in the larvae initially increased, followed by a decreasing trend. In a previous study, we found that *A. apis* inoculation of *A. cerana* larvae led to chalkbrood disease, reduced the larval survival rate, and significantly affected the activities of SOD, CAT, GST, and PPO [70]. This is indicative of the involvement of these four crucial antioxidant enzymes in the host response to *A. apis* infection. Here, nineteen DEcircRNAs, five DEmiRNAs, and three mRNAs shared by the 4-, 5-, and 6-day-old comparison groups were associated with three antioxidant enzymes of great importance, including SOD, CAT, and GST (Figure 7). This suggested that the DEcircRNA-mediated ceRNA network is potentially engaged in regulating the response of *A. c. cerana* worker larvae to *A. apis* infection. In the near future, functional dissection of key DEcircRNAs (e.g., RNAi via feeding siRNA targeting back-splicing sites) [71] and miRNAs (overexpression and knockdown via feeding the mimic and inhibitor) [72] will be conducted to explore the ceRNA metabolism underlying the *A. c. cerana* larval response to *A. apis* invasion.

When they are exposed to pathogens or parasites, insects initiate hemolymph-mediated cellular immune and fat-body-mediated humoral immune responses to defend against infection [73]. Phagocytosis and encapsulation are the classical insect cellular immune pathways, in which phagocytosis mainly engulfs pathogens with small molecular sizes, including bacteria and viruses, while pathogens with larger molecular sizes, such as parasites and nematodes, are removed by encapsulation and colonization [73]. Lemaitre and Hoffmann et al. [74] reported that large numbers of lamellocytes can be induced to differentiate from hemocyte precursors upon infection of larvae with parasitoid wasp eggs, forming a vesicular complex with melanism or producing ROS to engulf and kill the contents. Apoptosis, as a key component of the cellular immune system, is an active programmed cell death under polygenic control and plays an important role in the cellular response to infections by various pathogens. Zhang et al. [75] found that infection of *Spodopteralitura* by Autographa californica multicapsid nucleopolyhedro virus (AcMNPV) induced the activation of host cell apoptosis at the early stage to suppress viral propagation and spread. Here, cellular immune-relevant sub-networks were investigated, the results showed that 51 DEcircRNAs in the by 4-, 5-, and 6-day-old comparison groups potentially targeted 9 DEmiRNAs and further targeted 31 DEmRNAs. These target DEmRNAs were involved in six cellular immune pathways, such as apoptosis, melanogenesis, endocytosis, autophagy-animal, apoptosis - fly, and insect hormone biosynthesis (Figure 8A). Toll, IMD, and JAK/STAT are three vital signaling pathways, among which, the Toll signaling pathway mainly responds to Gram-positive and fungal infestations, while the IMD signaling pathway mainly responds to Gram-negative infestations [76]. The JAK/STAT signaling pathway is mainly involved in antiviral natural immunity and melanism [77]. It has been suggested that the Toll and IMD signaling pathways regulate insect gut flora and maintain gut immune homeostasis [78, 79]. Both the Toll and IMD signaling pathways result in the activation of Nf-*κ*b transcription factors and translocation into the nucleus, where they up-regulate the expression of genes encoding AMPs and other genes [80]. Lin et al. [81] analyzed the expression profiles of immune genes in the fourth-instar *Plutella xylostella* larval midgut infected by *Staphylococcus aureus, Escherichia coli*, or *Pichia pastoris*. They found that the expression levels of immune genes related to the Toll, IMD, and JAK/STAT signaling pathways were significantly up-regulated, which is suggestive of the participation of these three immune pathways in host immunity. Sun et al. [79] detected that the Toll receptor genes *Toll 1A* and *Toll 5A* in *Anopheles stephensi* were highly activated in response to *Beauveria bassiana* infection, *Toll 1A* and *Toll 5A* regulated the homeostasis of gut microbiota. After suppressing the JAK-STAT signaling pathway by silencing the negative regulator PIAS in *Aedes aegypti* based on RNAi, Souza-Neto et al. [82] observed that the expression levels of one putative Toll-receptor-associated gene was up-regulated and the host susceptibility to dengue virus infection significant decreased. In this study, target DEmRNAs were engaged in six humoral immune pathways, including the Toll and Imd, Toll-like receptor, NF-kappa B, Jak-STAT, and MAPK signaling pathways (Figure 8B). In summary, these results demonstrated that corresponding DEcircRNAs and their involved ceRNA network were likely to regulate the expression of the aforementioned immune-pathway-related genes, followed by participation in the response of *A. c. cerana* worker larvae to *A. apis* infection.

Unlike linear RNAs, circRNAs lack the 5′ cap and the poly(A) tail; hence, they cannot encode proteins through classical translation mechanisms. However, recent studies have suggested that some circRNAs including IRES elements and ORFs are able to encode small peptides or proteins with a biological function [83]. Zhang et al. [84] identified that vSP27, translated from a BmCPV-derived circular RNA, induced the generation of ROS and activated the NF-κB signaling pathway, induced the expression of antimicrobial peptides, and suppressed BmCPV infection. In silkworms, circEgg encoded circEgg-P122, a protein with 122 amino acid residues, and inhibited tri-methylation of histone H3 and lysine 9 [85]. In this current study, 27, 26, and 24 IRES as well as 61, 52, and 40 ORFs were identified in DEcircRNAs in the 4-, 5-, and 6-day-old comparison groups, respectively. This implied that these DEcircRNAs had protein-encoding potential. Additionally, the ORFs within DEcircRNAs could be annotated to the cAMP signaling pathway, metabolic pathways, and ABC transporters, in addition to several immune pathways such as the insulin signaling pathway, melanogenesis, phagosomes, lysosomes, the MAPK signaling pathway (fly), apoptosis (fly), and endocytosis. Together, these results indicate that some DEcircRNAs may modulate many aspects during the larval response to *A. apis* infection by encoding related proteins.

## 5. Conclusions

In conclusion, 3178 circRNAs were identified in the *A. c. cerana* worker larval guts. The expression pattern of circRNAs in the larval guts was changed due to *A. apis* invasion; corresponding DEcircRNAs were potentially engaged in host responses including the immune response to *A. apis* infection through versatile mechanisms (Figure 11), such as the regulation of the transcription of parental genes, absorption of target miRNAs via ceRNA networks, and translation into proteins. The findings from this study lay a foundation for further investigation of the function and mechanism of circRNAs regulating the host *A. apis* response, provide novel and valuable insights into the interactions between *A. cerana* larvae and *A. apis*, and offer candidate biomarkers and molecular targets for the diagnosis and control of chalkbrood disease.

**Figure 11.**
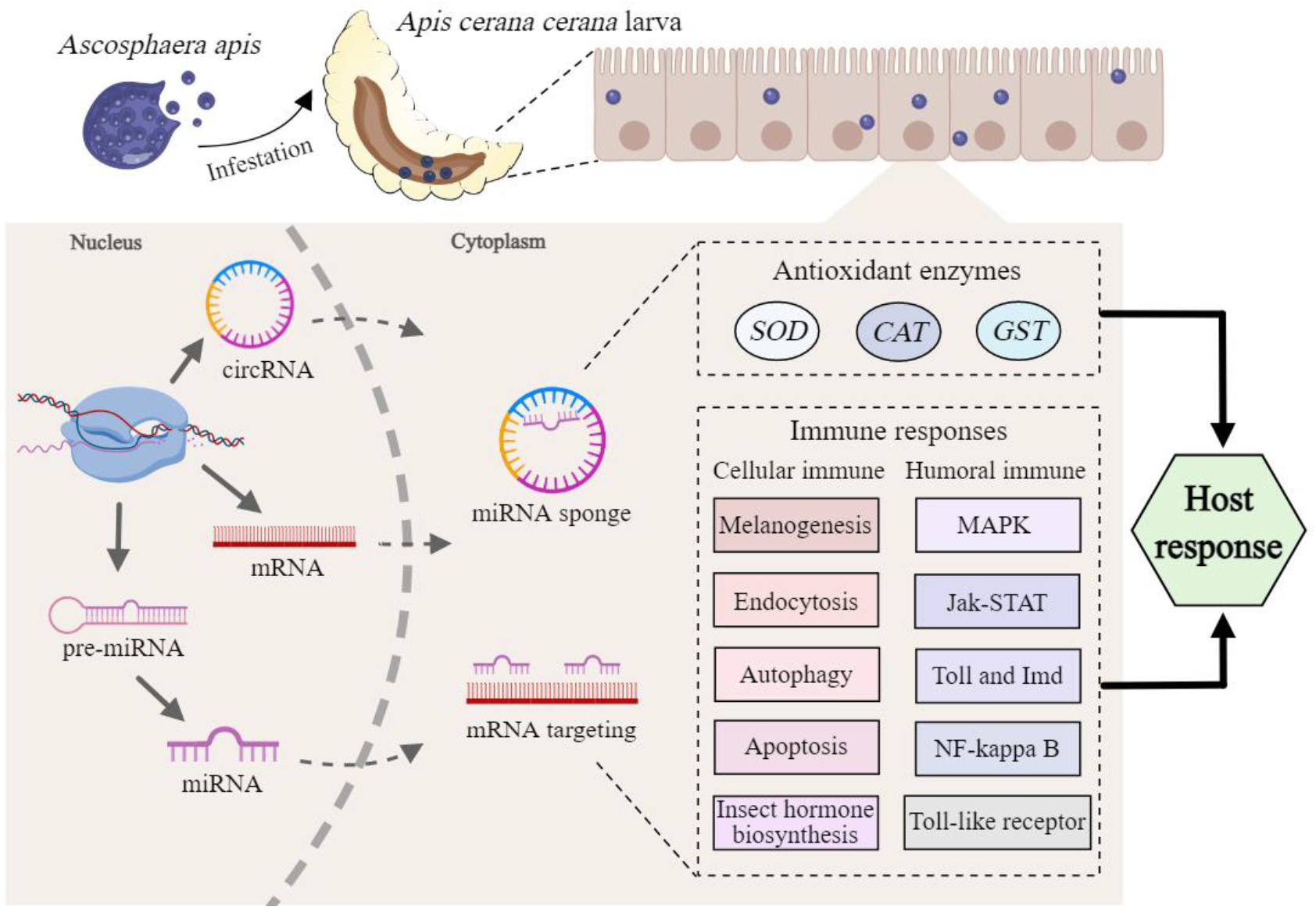
A hypothetical schematic diagram of circRNA-mediated immune responses of *A. c. cerana larvae* to *A. apis* invasion. This diagram was created with MedPeer website (www.medpeer.cn).

## Author Contributions

R.G. and D.C. designed this research; R.G., K.Z. contributed to the writing of the article; K.Z., X.G., X.J., S.G., H.Z., Y.S., K.L., P.Z., and M.C. conducted experiments and data analyses; Y.W., and Z.H. supervision; R.G., Z.F., and D.C. funding acquisition. All authors have read and agreed to the published version of the manuscript.

## Institutional Review Board Statement

Not applicable.

## Informed Consent Statement

Informed consent was obtained from all subjects involved in the study.

## Data Availability Statement

All the data is contained within the article.

## Acknowledgments

All of the authors thank reviewers and editors for their valuable comments and recommendations. R.G. appreciates his beloved wife and daughter for their great love and assistance.

